# Learning to Adapt - Deep Reinforcement Learning in Treatment-Resistant Prostate Cancer

**DOI:** 10.1101/2023.04.28.538766

**Authors:** Kit Gallagher, Maximillian Strobl, Robert Gatenby, Philip Maini, Alexander Anderson

## Abstract

Standard-of-care treatment regimes have long been designed to for maximal cell kill, yet these strategies often fail when applied to treatment–resistant tumors, resulting in patient relapse. Adaptive treatment strategies have been developed as an alternative approach, harnessing intra-tumoral competition to suppress the growth of treatment resistant populations, to delay or even prevent tumor progression. Following recent clinical implementations of adaptive therapy, it is of significant interest to optimise adaptive treatment protocols. We propose the application of deep reinforcement learning models to provide generalised solutions within adaptive drug scheduling, and demonstrate this framework can outperform the current adaptive protocols, extending time to progression by up to a quarter. This strategy is robust to varying model parameterisations, and the underlying tumor model. We demonstrate the deep learning framework can produce interpretable, adaptive strategies based on a single tumor burden threshold, replicating and informing a novel, analytically–derived optimal treatment strategy with no knowledge of the underlying mathematical tumor model. This approach is highly relevant beyond the simple, analytically–tractable tumor model considered here, demonstrating the capability of deep learning frameworks to help inform and develop treatment strategies in complex settings. Finally, we propose a pathway to integrate mechanistic modelling with DRL to tailor generalist treatment strategies to individual patients in the clinic, generating personalised treatment schedules that consistently outperform clinical standard-of-care protocols.

## 1 Introduction

Drug resistance is responsible for up to 90% of cancer-related deaths [1]. It can be present before treatment (intrinsic) or emerge during therapy (acquired) and is driven by a combination of genetic, epi-genetic and environmental processes [2]. Much of modern cancer research has focused on developing novel therapies to overcome resistance but, especially in the metastatic setting, cure rates remain low and all too often benefits are short-lived (e.g. [3, 4]).

Conventional, standard-of-care treatment schedules in chemotherapy are based on the maximum tolerated dose (MTD) principle. This argues for the administration of treatment at as high a dose and frequency as tolerable, in order to maximize cell kill and thereby the chance of cure [5]. However, over the past years it has become increasingly clear that cancers, in particular metastatic cancers, are complex and spatio-temporally heterogeneous and actively evolve under treatment [6]. This has prompted a re-thinking of the MTD paradigm, and has stimulated a growing body of research demonstrating that changes in drug-scheduling could delay drug resistance [7, 8, 9, 10]. One particularly promising approach is so-called “adaptive therapy” (AT) which is based on the principle of “competitive control”, wherein it is hypothesized that while resistant cells, or their pre-cursors, may exist prior to treatment, their expansion is restricted by competition for space and resources with the drug-sensitive subpopulation [9, 11]. MTD treatment, on the other hand, facilitates competitive release since it rapidly depletes sensitive cells and the associated competitive suppression, which allows expansion of the resistant subpopulation and causes progression [12, 13]. To address this, Gatenby et al. [9, 11] proposed AT, which dynamically modulates treatment, so as to maintain a pool of drug-sensitive cells that compete with emerging resistance, seeking to control the tumor rather than eliminating it. These adaptive strategies exploit both spatial and resource competition between drug-sensitive and -resistant cells [14, 15, 16], and the fitness costs associated with drug resistance [17, 18]. AT has been shown to extend the time to progression (TTP) *in vivo* for breast cancer [19], ovarian [9] and lung cancer [20], and melanoma [21] and has, most recently, delivered promising results in a pilot clinical trial in metastatic castrate-resistant prostate cancer [22, 23].

Castrate-resistant prostate cancer is treated with androgen deprivation therapy, such as abiraterone which inhibits CYP17A, an enzyme for testosterone auto-production [24]. Standard of care is given at MTD via continuous administration until radiographic progression, which occurs after a median of 16.5 months [25]. Zhang et al. [22, 23] instead applied an adaptive strategy (AT50) where an identical dose was given until the tumor burden had reduced by 50% relative to baseline and was subsequently withheld until it returned to baseline (NCT02415621). To monitor burden, they combined radiographic imaging with measurements of Prostate Specific Antigen (PSA) levels, an established serum biomarker [26, 27], which allowed for more frequent, monthly tracking of the disease. In comparison with a matched historical control receiving continuous dosing, the study found that patients undergoing AT had a 19.2 month increase in median progression-free survival, while receiving 46% less drug on average [23].

Although these results are promising, they also highlight the need for further research into how exactly we adapt therapy: only 4 of 17 patients on the trial were able to achieve long term disease control [23], and there was significant variation in the adaptive cycling dynamics between patients (exemplified in Figure 1a). Previously we, and others [17, 28, 29, 30], have investigated how the threshold of tumor burden at which treatment is withdrawn in the AT50 protocol affects outcome. This showed that increasing the threshold, so that treatment is withdrawn earlier and at a higher average tumor burden, increases competitive suppression and thereby TTP. Consequently, it has been proposed [29, 28] that the tumor could even be allowed to increase in size beyond its baseline level in what Brady et al [31] have called “range-adaptive” AT. At the same time, these benefits are subject to a trade-off: a higher tumor burden also means an increased risks of phenotype switching, *de novo* mutations, and metastasis [10, 29, 30], indicating that the question of when and how to adapt therapy should ideally be answered on a personalized basis. Furthermore, while the AT50 protocol represents an important first step, it is limited in its generalisability. How would we, for example, integrate multiple drugs, unforeseen treatment interruptions (e.g. for delays in data acquisition or toxicity), or patient-specific treatment goals?

**Figure 1:**
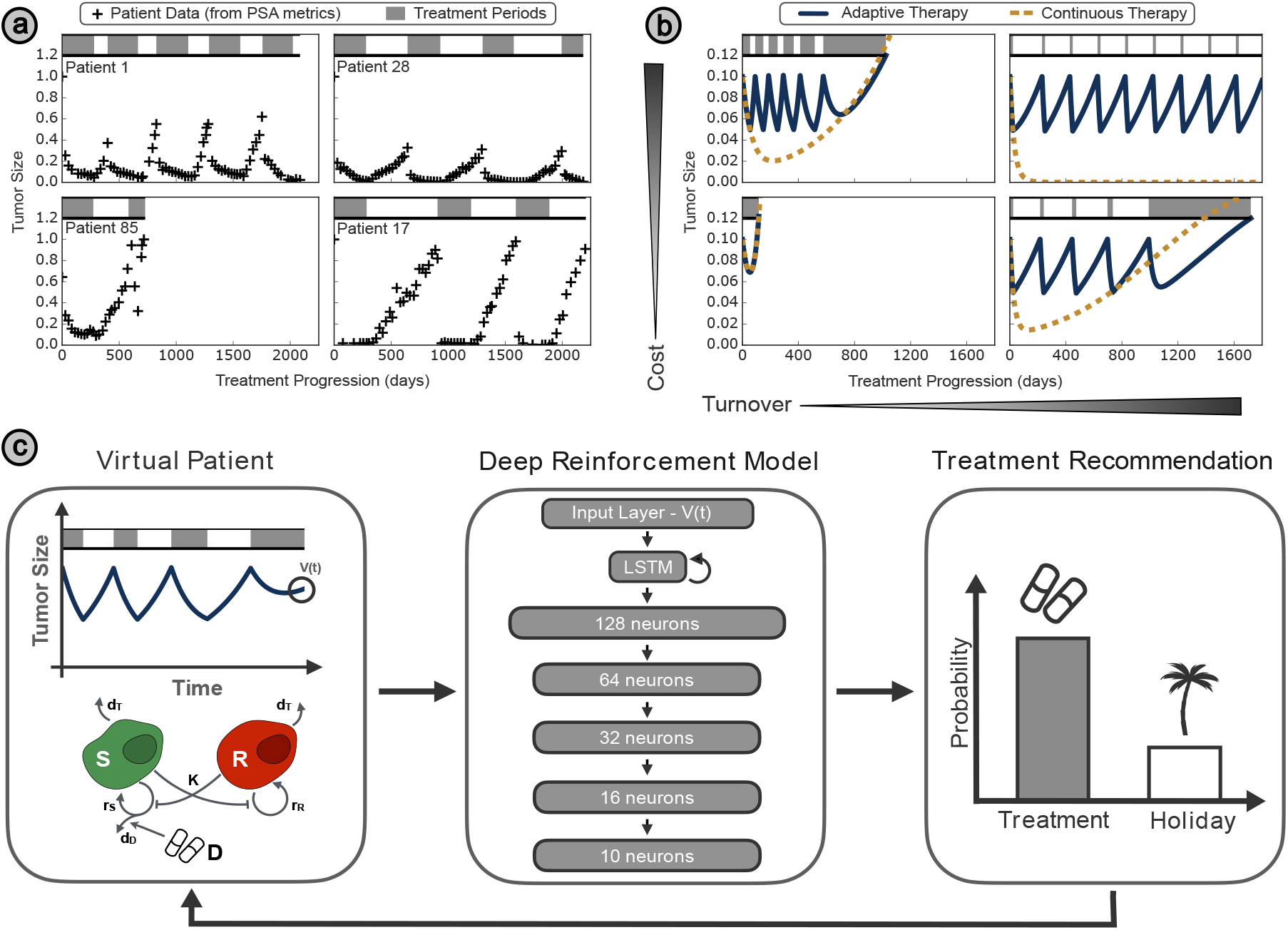
**(a)** Example treatment records of patients from Zhang et al. [22] demonstrating the differing treatment responses observed under this model. Tumour size is quantified by levels of PSA in the blood, and normalised relative to the initial value. The treatment schedule is given by the bar at the top of each plot, with shaded regions corresponding to treatment periods, separated by treatment holidays in white. **(b)** The variety in patient response to treatment can be represented by a mathematical tumor model, varying the cellular turnover and cost associated with drug resistance. **(c)** Tumour metrics generated by the virtual patient (full details given in Section 2.1) fed into a deep learning framework, which returns a treatment recommendation. A random treatment decision is sampled based on this, and then propagated back to the virtual patient, which simulates the outcome of this decision over the next timestep.

From chat bots to self-driving cars, deep learning techniques are revolutionizing the world around us. These ‘deep’ methods use artificial neural networks with many intricately connected layers, enabling them to learn highly complex relationships between input variables [32]. While initially focused on classification problems, such as cancer diagnosis [33], most recently so-called Deep Reinforcement Learning (DRL) has extended these methods to decision-making in dynamic and complex environments such as the board game Go [34], autonomous vehicles [35], or problems in healthcare [36].

DRL frameworks have achieved success in a range of drug scheduling problems, ranging from immune response after transplant surgery [37] to controlling drug resistance in bacteria [38, 39]. To do so, at each timestep a deep learning agent is given information on the state of the system (e.g. tumor size) and its output is used to choose from a set of possible actions (e.g. treat vs not treat) [40]. To learn its strategy, the agent is trained through a process of trial and error to maximize a reward function that remunerates success (e.g. tumor shrinkage or cure), and penalises unfavorable events (e.g. excess drug toxicity) [41]. DRL schemes are particularly well suited to this task, as they may account for the long term effects of actions when maximising outcomes, even when the relationship between actions and outcomes is not fully known [42]. For example, using absolute neutrophil counts as a biomarker for chemotherapy-associated toxicity, Maier et al. [43] proposed a framework for adjusting subsequent drug doses, to reduce toxicity in cancer patients. They demonstrated that reinforcement learning frameworks have the potential to substantially reduce the incidence of neutropenia, and provide insight into the patient factors that determine treatment recommendations.

In another example, Engelhardt [38] developed a DRL framework for the analogous problem of antibiotic resistance, to predict precision dosing that adaptively targets harmful cell populations with variable drug susceptibility and resistance levels. They introduce a simple DRL framework capable of suppressing cell proliferation and demonstrate robustness to changes in model parameters, based on discrete-time feedback on the targeted cell population structure. Given the context of antibiotic resistance, their model is based on the assumption that all strains have some degree of drug response, and hence can ultimately be eliminated, which differs significantly from the reality of non treatment-responsive tumors in cancer treatment. Furthermore, Eastman et al. [44] have also shown that chemotherapeutic schedules derived on-the-fly (through reinforcement learning) are more robust to changes in the tumor behaviour than strategies determined a priori through global optimisation (such as via classical optimal control methods).

However, while DRL methods are very promising, their translation into clinical practice faces a key challenge: unlike a game of chess, a patient’s treatment plan cannot be replayed until the DRL agent has learned its strategy. To address this, studies to-date have used mathematical models to serve as “virtual patients”, generating the vast quantities of data needed to train a machine learning algorithm and predicting how the tumor may respond to any hypothetical treatment scenario that could not easily be tested in the clinic. Yet this inherently links the so-learned strategies to the assumptions of the specific, underlying model. If those assumptions are not met, then how robust are DRL methods when a patient’s disease behaves differently to the training model? Can we adjust the method to learn while we are treating? And how does this compare to standard-of-care treatment techniques?

The aim of this paper is to investigate whether deep learning techniques may allow us to integrate evolutionary principles and mathematical models more directly into AT decision-making and uncover novel, clinically-translatable AT approaches. Using a previously characterized and calibrated Lotka–Volterra mathematical model to simulate the intra-tumor ecological dynamics (see Figure 1b) we test the ability of a DRL algorithm to guide therapy (Figure 1c). We demonstrate that this framework can outperform both standard of care and conventional adaptive strategies, and discuss how it can help to uncover interpretable and rational principles for optimal scheduling design. In the second part of this paper, we turn to the key question of how to make DRL-based scheduling clinically feasible, when we cannot be certain about the specific characteristics of a patient’s disease, and there is no way to replay or take back a treatment decision once it is made. We apply our framework to a virtual patient cohort with a range of characteristics, demonstrating its robustness to certain changes in tumor parameters and dynamics. To conclude, we propose a framework in which we integrate mechanistic mathematical models with deep reinforcement learning to deliver dynamic, patient-specific treatment scheduling.

## 2 Methods

### 2.1 Virtual Patient Model

To rapidly and safely benchmark DRL-informed AT, we use a mathematical model to simulate the treatment response of a “virtual patient”. We adopt the 2-population Lotka–Volterra model introduced by Strobl et al. [17], where *S*(*t*) is the number of sensitive cells, and *R*(*t*) is the number of resistant cells:

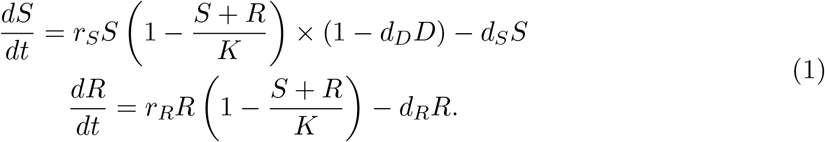

Briefly, the model assumes sensitive and resistant cells proliferate and die at rates *r*_*S*_ and *r*_*R*_, and *d*_*S*_ and *d*_*R*_, respectively, and compete for a shared carrying capacity, *K*. Treatment is assumed to kill sensitive cells at a rate that is proportional to the population’s growth rate and the drug concentration, *D*(*t*). Resistant cells are assumed to be fully resistant and can be subject to a cost, so that, *r*_*S*_ *≥ r*_*R*_.

To simplify notation we define the “cost of resistance” as defined by the difference between resistant and sensitive cell growth rates (1 *— r*_*R*_*/r*_*S*_). Similarly, we define cell turnover as the relative proliferation and death rates of sensitive cells *d*_*S*_*/r*_*S*_. Parameter values were adopted from Strobl et al. [17] and are given in Table 1 from Supplementary Information 2.

**Table 1:** Terms, and their default values, in the default reward function for the DRL framework.

| Name | Value | Circumstance | Motivation |
| --- | --- | --- | --- |
| Base | 0.1 | Per timestep survived | Maximises TTP |
| Holiday | 0.05 | Per timestep without treatment | Encourages intermittent therapy |
| Progression | - 0.1 | Upon progression (20% growth from $N_0$) | Punishes progression |
| Survival | 5 | Upon survival for 30 years | Reward indefinite survival |

### 2.2 Adaptive Therapy

We benchmark DRL-informed AT against the following two protocols:

1. **CT** – Continuous Therapy (Standard of Care):

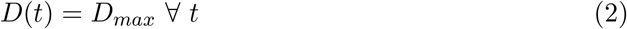

2 **AT50** – AT schedule used in the pilot clinical trial by Zhang et al. [22, 23]. Treatment is given until a 50% decrease from initial size (*N*_0_) is achieved, then withdrawn until tumor returns to itsinitial size:

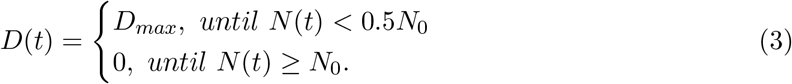

We compare schedules based on their TTP, where we define progression as a 20% growth from the initial size, as done in prior studies in this area (e.g. [14, 17, 29]).

### 2.3 Deep Learning Model

To test the feasibility and potential benefits of DRL-driven AT, we develop a prototype in which we use the asynchronous, advantage Actor–Critic (A3C) network pioneered by Mnih et al. [45] to drive treatment decision-making. The A3C framework consists of a global network, with many duplicates (workers) updating to it asynchronously during training (Figure 2), which avoids the high computational costs and specialist architecture requirements associated with GPU-based deep-learning algorithms [46]. The network receives as input the current tumor size and outputs a policy score for each of the two possible actions (treat vs drug-holiday) which reflects which is predicted to be the more successful. To decide whether or not to treat in the next time step, the scores are converted into probabilities, and an action is chosen probabilistically. The network architecture is depicted in Figure 1c, and further details as well as a pseudocode representation are given in the Supplementary Information 1.

**Figure 2:**
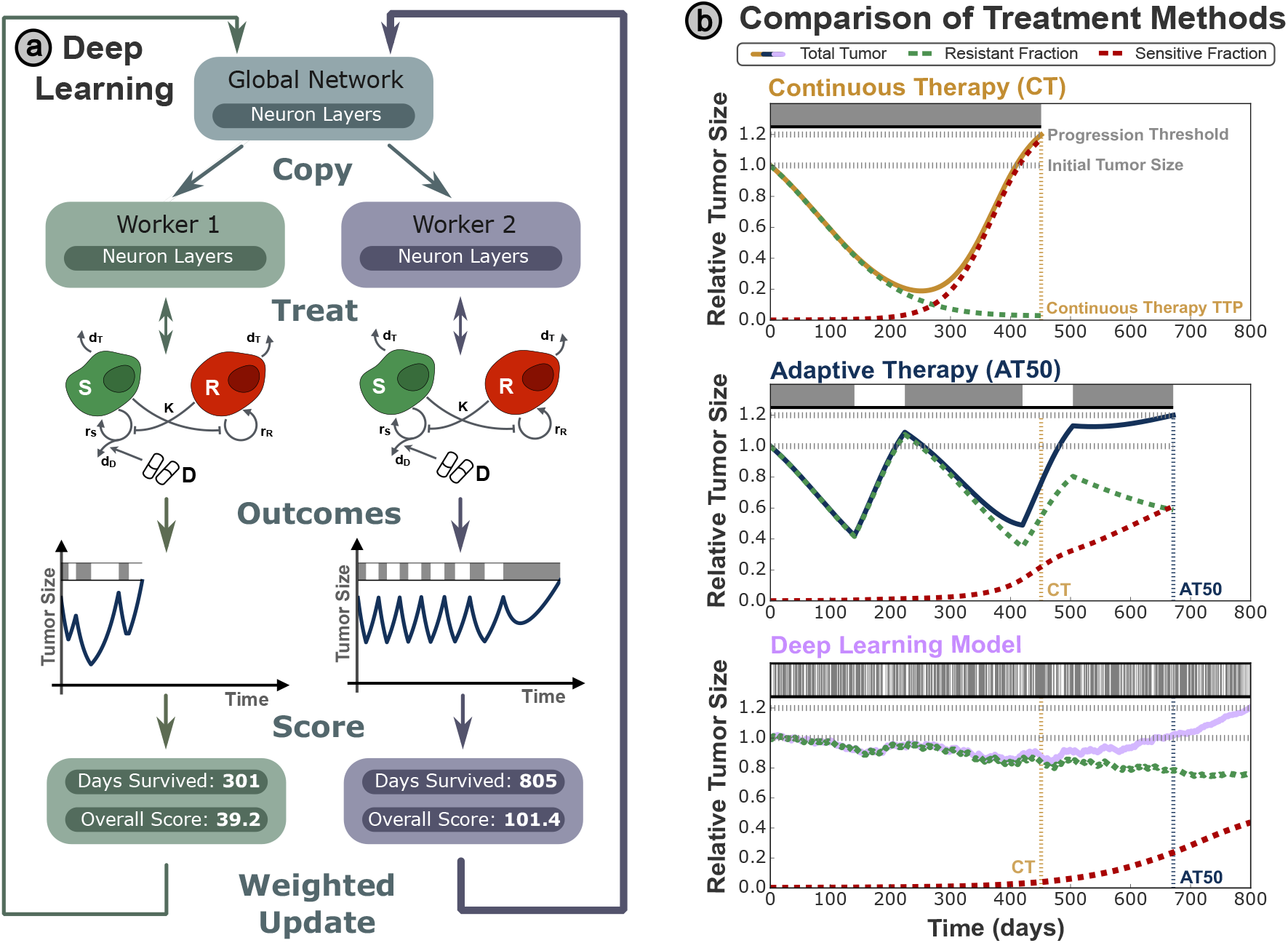
**(a)** The training process, where treatment strategies are optimised by comparing perturbed copies of the DRL network evaluated on the virtual patient system, and adjusting the global DRL network based on the relative performance of each copy. **(b)** A comparison of different treatment strategies, with the treatment schedule recorded in a bar at the top of each panel, where shaded regions correspond to drug treatment. The tumor profile is shown over time, with red and green lines depicting, respectively, the resistant and sensitive tumor sub-populations. Treatment terminates when the total tumor size reaches 120% of the initial size, where the tumor is deemed to have reached progression.

Because the decision-making in the DRL framework is probabilistic, the outcome can vary between iterations. To account for this, we report results as averages over 100 evaluations (unless otherwise stated). This number was determined to be sufficient based on a consistency analysis (not shown).

### 2.4 Reward Function

The DRL framework learns treatment strategies through optimising an objective function, which returns a reward calculated from the most recent state and treatment choice at each timestep. This reward is based the number of timesteps survived, encouraging the model to maximise TTP, as well as a bonus for treatment holidays to incentivise intermittent treatment strategies. Rewards associated with the final size of the tumor are also used to highlight actions strongly linked with positive and negative final outcomes. A list of all terms, alongside their default values and the circumstances under which they are rewarded, is given in Table 1.

Discounting is applied to the reward function to determine how important future rewards are to the current state, where a reward that occurs *N* steps in the future is multiplied by *γ*^*N*^, for discounting factor *γ ∈* (0, 1). As well as ensuring convergence of the reward sum, this factor is used to tune prioritisation of short and long timescales in the reward function. A value very close to unity (*γ* = 0.999) is used throughout to prioritise overall outcomes over shorter-term benefits.

### 2.5 Data Availability

All methods, along with the DRL framework, will be made available upon publication. Clinical data from Bruchovsky et al. [47] was obtained from https://www.nicholasbruchovsky.com.

## 3 Results

The aim of this paper is to investigate whether deep learning can inform cancer therapy scheduling and do so in a clinically translatable fashion. To investigate this, we tested DRL-guided adaptive therapy on cohorts of virtual patients in which the tumor dynamics were driven by a previously established mathematical model [17]. This model assumes that the tumor is composed of treatment-sensitive and resistant cells and that these compete with each other for resources in a Lotka–Volterra fashion (Figure 2).

### 3.1 DRL-guided adaptive therapy can outperform current clinical strategies

As a first step, we carried out a proof-of-principle case study on a patient representing an initially responsive, but relatively rapidly progressing, disease (Figure 2; parameters taken from Strobl et al [17]). Akin to the AT50 algorithm by Zhang et al [22], in our DRL-guided protocol, the patient’s tumor burden is monitored and the decision of whether or not to continue to treat is updated at a fixed “decision-frequency”. Importantly, however, this decision is not based on a fixed rule-of-thumb, but on a DRL algorithm which is carefully trained prior to deployment. During this training process, the DRL framework is applied to a cohort of “training patients”, its performance is scored according to the reward function, and the parameters of the underlying neural network are refined until it converges on a final decision-making policy (Figure 2a).

To test whether the DRL could learn an effective treatment strategy, we initially made the idealised assumption that we could train on a patient identical to the one on which the framework will be deployed. Following 2600 training epochs, we find that DRL-guided treatment is able to control the tumor for longer (Figure 2b). Through dynamic adjustment of treatment it achieves an average TTP of 745 days over 100 independent simulations (95% confidence interval:[688d, 802d]). The variability in performance is due to the stochastic nature of the DRL decisionmaking (see Supplementary Information 5.1 for examples). In comparison, CT and the AT50 therapy progress after 450 and 662 days, respectively (Figure 2b). We conclude that DRL-guided therapy is, in principle, feasible and can improve upon the current AT50 rule-of-thumb approach.

### 3.2 Reducing decision-making frequency can increase performance

To investigate whether, and how, this framework could be employed in practice, we next analysed sensitivity to key parameters in the training and deployment process. In the previous section, we made the somewhat unrealistic assumption that the DRL framework receives tumor size input and re-evaluates the treatment strategy on a daily basis. Clearly this would be both difficult and costly to implement clinically. Interestingly, and somewhat counter-intuitively, we found that reducing the treatment frequency of the model increases the expected TTP, despite the reduced information and intervention frequency (Figure 3a). In addition, this reduces the computational cost per patient during training. We hypothesize that this improvement in performance is because less frequent decisions are more impactful, enabling the DRL framework to better separate the meaningful trends in the underlying tumor dynamics from random noise in the decision process. This is reflected in the reduced variation in TTP for longer treatment frequencies (Figure 3b-c) and is consistent across different timescales, as shown in Table 2.

**Table 2:** The time to progression (TTP) for three DRL frameworks trained with different decision making frequencies, each one evaluated with varying decision-making frequencies. Premature failure may occur when evaluated on timescales greater then that used for training. For comparison, the TTP under the AT50 scheme is 672 days, and 451 days under conventional CT.

| TTP (days) | Training Decision Frequency |  |  |  |
| --- | --- | --- | --- | --- |
| Evaluation Decision Frequency |  | Daily | Weekly | Monthly |
|  | Daily | 745 | 1141 | 1281 |
|  | Weekly | 571 | 1116 | 1341 |
|  | Monthly | 414 | 353 | 1388 |

**Figure 3:**
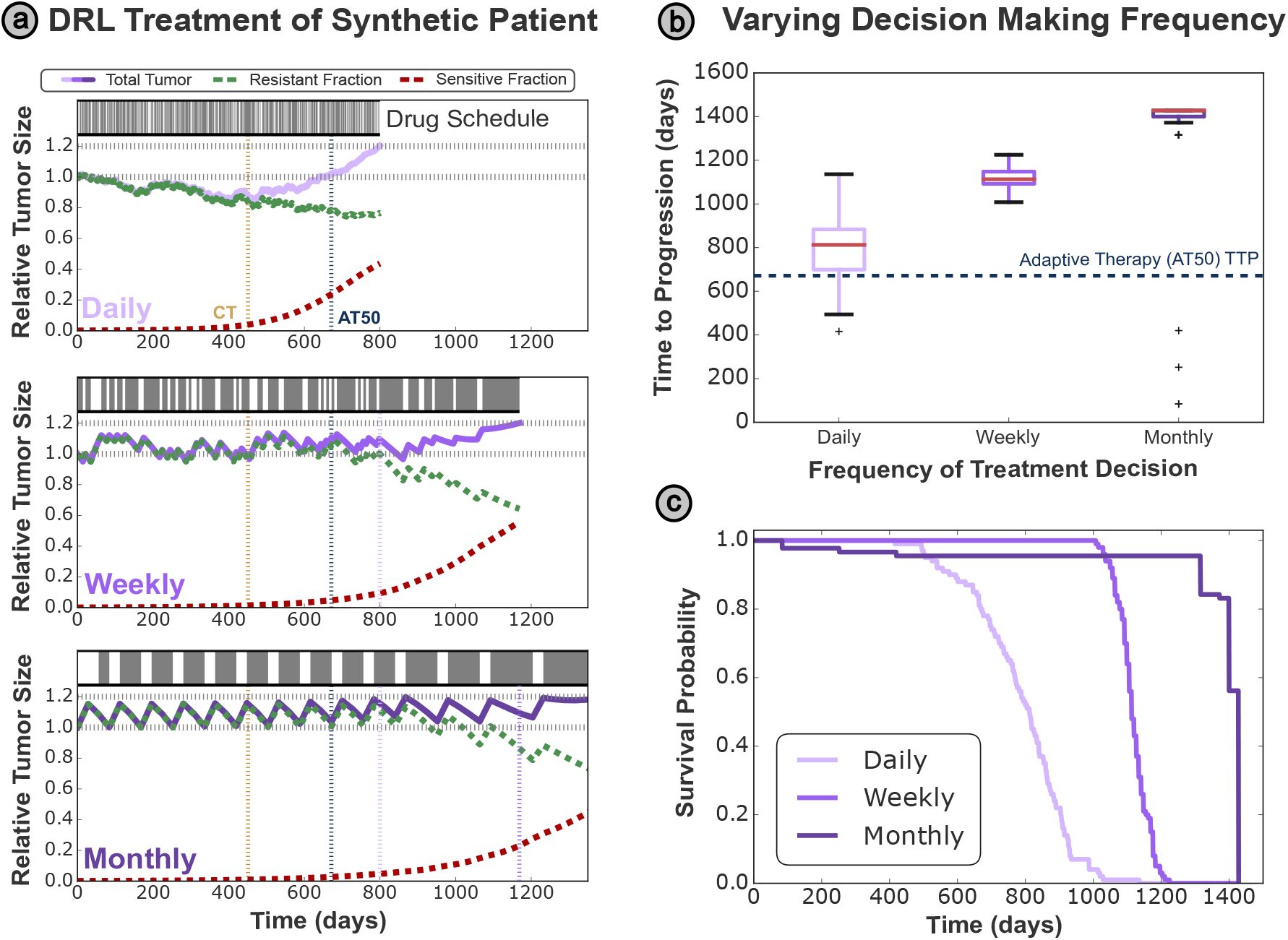
**(a)** DRL framework performance for different treatment intervals after training on a single patient profile. Within this range, longer treatment intervals in DRL training correspond to greater TTP values and more consistent treatment schedules. **(b)** The expected TTP from the DRL framework under different treatment frequencies, in comparison to the expected value from AT50. There is distinct separation between the interquartile ranges for each decision frequency, with the variation in TTP quantified by error bars representing two standard deviations. **(c)** The relative improvements of treatment strategies with less frequent decision making, across a cohort of 100 virtual patients.

We also considered the role of the treatment-frequency during evaluation. In this case, we find no benefit from reducing the treatment interval used in training, as the model has been optimised for a specific treatment schedule (shown above the diagonal in Table 2). However, if the treatment interval is increased in evaluation compred to training, TTP is significantly reduced. This is because the DRL framework underestimates the extent of growth during each interval and fails to apply sufficient treatment. The tumor then grows above the threshold for progression even though it is still sensitive and could have been kept in check with further treatment, a phenomenon we will refer to as “premature progression” (Supplementary Figure 1c). To sum up, choosing the frequency at which to consult the DRL framework will require a careful balance between leaving sufficient time to learn from past decisions, whilst also providing frequent enough decisions to react to the tumor’s response, or lack thereof.

### 3.3 DRL frameworks can replicate optimal analytic strategies

While the performance gain of out ‘black box’ DRL framework seems impressive, in order for this approach to be translated to clinic we need to understand why it does what it does. A clear interpretation of the DRL decision framework will make it more transparent and more believable. To do this, we will convert the ‘treatment probability’ output from the DRL network into a treatment recommendation where higher probabilities correspond to a greater certainty that the patient should receive treatment for the next time interval. Investigating the relationship between the current tumor size (the input to the network) and the treatment probability (the output from the network), we find that the DRL framework has learnt a strongly defined sigmoidal relationship (as depicted in Figure 4a). We can interpret this as a simple binary prediction on the best treatment course dependent on the tumor size; when the current tumor size is above the critical size *N* ^*\**^ we should provide treatment, whereas if the tumor size is below this critical size then the patient would benefit more from a treatment holiday. Such a strategy could therefore be easily implemented in the clinic, with this critical size determined by the DRL network and personalised to each patients tumor burden.

**Figure 4:**
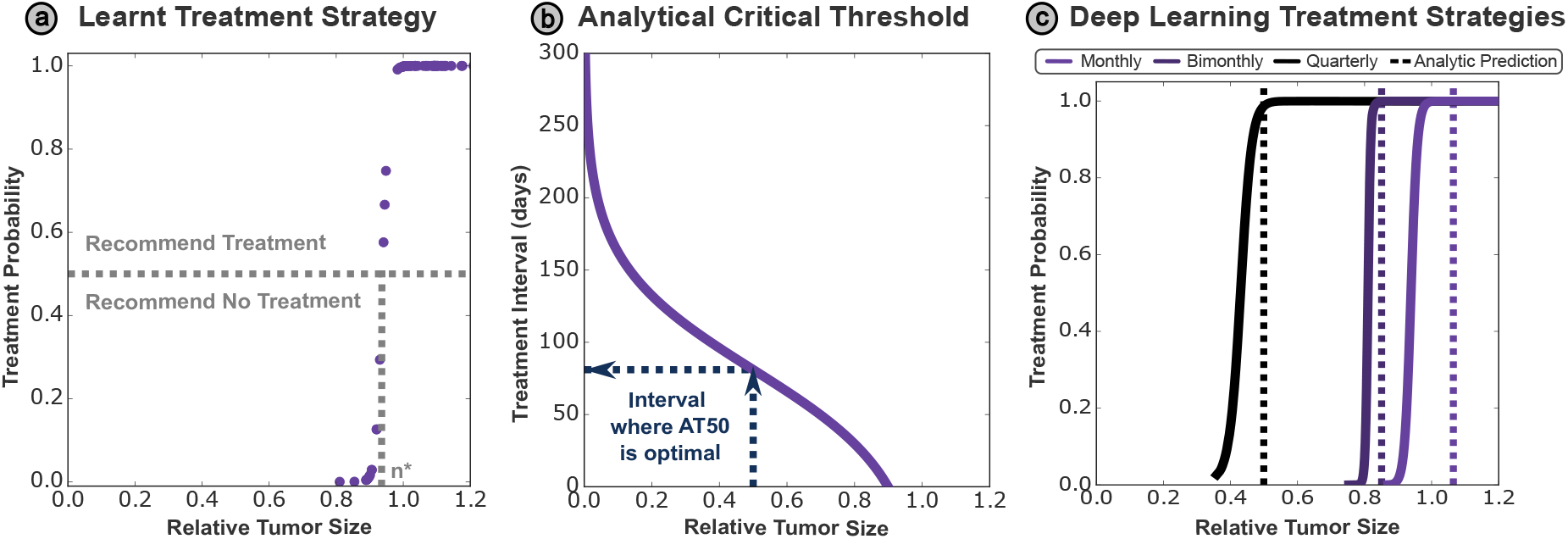
**(a)** The DRL learns a well-defined sigmoidal strategy for recommending treatment based on the current tumor size, with treatment almost certain over a threshold size (0.83*n*_0_). **(b)** The optimal threshold interval depends on the treatment threshold used - AT50 is therefore only optimal for one specific treatment interval (81 days). If the treatment interval can be reduced from this, then AT50 is suboptimal, while if it must be less frequent, then AT50 will fail prematurely. **(c)** The DRL is able to replicate threshold size (above which treatment should be provided) predicted by the analytic model for a range of treatment frequencies.

We can think about this critical size, for a given patient, as the largest tumor size where it is still safe to give a treatment holiday - above this size it is likely that their tumor will reach progression in the next time interval, undergoing premature failure. However, the DRL framework has learnt to keep the patient’s tumor as close to this critical size as possible, since a larger sensitive sub-population of cells will result in greater suppression of the drug-resistant subpopulation, delaying the patient’s progression. In this way, the DRL framework replicates basic strategies known to be analytically optimal, from the field of Optimal Control Theory [10, 29]. This critical size is inherently dependent on the treatment interval, and how fast the tumor can grow within this fixed period of time. While patients may have different tumor dynamics, the treatment interval is typically the same for all patients, due practical constraints within the clinic. This would mean that different patients could respond better to different threshold sizes for treatment, and the “one size fits all” approach of AT50 is sub-optimal for most patients. If treatment is given below this threshold then we suppress the sensitive subpopulation within the tumor unnecessarily, while if treatment is withheld above this threshold then we risk the patient undergoing premature progression before the next treatment decision.

For this simple model (1), we can derive a mathematical expression for the optimal treatment threshold based on a pre-determined treatment interval */tau*. This allows us to calculate the exact treatment threshold that will be optimal for each individual patient, based on their tumor parameters as defined in (1) and how frequently they should be treated. The derivation is outlined in the Supplementary Information, with the final expression for the critical treatment threshold *N* ^*\**^ given as:

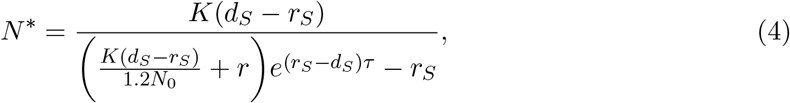

and plotted in Figure 4b. This demonstrates the extent of the variation in optimal threshold for a single patient depending on how frequently treatment decisions may be made. Given the treatment threshold in AT50 is fixed at 50%, there is only a single treatment interval (81 days) where AT50 is optimal - more frequent treatment re-evaluation has no additional benefit and so the patient progresses sooner than they would under an optimal treatment scheme. Conversely, premature progression can also occur in conventional AT when treatment is reevaluated insufficiently frequently, an example of which is given in Supplementary Figure 1a.

To investigate whether the DRL could replicate this analytic result, we retrained the DRL framework at three different treatment frequencies, plotting the treatment curves for each in Figure 4c. These show good agreement with the analytically derived values, demonstrating that the treatment strategy learnt by the DRL framework is close to optimal. We notice that the treatment thresholds of the DRL are typically below the theoretical value; this slightly conservative approach enables the framework to account for the inherent stochasticity in the DRL decision making process. This accuracy across a range of treatment frequencies is remarkable, as the mathematical intricacies of the Lotka–Volterra model are completely hidden from the DRL framework. This highlights a significant advantage of the DRL framework over analytic methods, since it was able to replicate complex analytic expressions numerically with no explicit knowledge of the underlying mathematical model or its solution.

To summarise, we have demonstrated how to identify clinically-actionable strategies from a DRL network, giving practitioners confidence in the treatment decisions it recommends. We have also demonstrated that these strategies represent optimal protocols (which may in this simple case be derived analytically), giving confidence that the DRL may be applied to more complex treatment paradigms (with multiple drugs or dosing levels) where mathematical methods cannot readily derive an optimal treatment schedule.

### 3.4 DRL frameworks can be robust to variation in patient parameters

Given this success in training the DRL framework on a single patient profile, we now look to extend that to encompass the broad spectrum of patients that are seen within the clinic. Now that we have shown that the critical treatment threshold will vary between patients, any DRL with clinical relevance must be robust to variation in the underlying tumor dynamics. As well as accounting for variation between patients, this also accounts for our imperfect knowledge of each patient’s underlying parameters, obtained by fitting to noisy clinical data.

To replicate tumor dynamics observed in the clinic, we consider patient data from a prospective Phase II trial of intermittent androgen suppression for locally advanced prostate cancer, conducted by Bruchovsky et al. [47]. This trial used an intermittent strategy based on adaptive therapy to treat biochemical recurrence after radiotherapy, with a continuous lead-in treatment period before allocating treatment holidays according to patient PSA metrics. We formed a cohort of virtual patients from these data, comprised of 67 patients in this trial that did not develop a metastasis. The behavior of these virtual patients is determined by the Lotka–Volterra model introduced in section 2.1, with parameter sets as determined by individually fitting to each patient’s treatment history (Strobl et al. [17]).

Simulating these parameter sets with the virtual patient model (1), however, reveals that only seven out of the 67 patients are expected to reach progression within 5000 days. This raises a small discrepancy with the original clinical data, where 12 of the patients were marked as reaching progression during the six year study period. This can be attributed to a differing progression criteria used by Bruchovsky et al. [47], based on both PSA and testosterone rises under treatment and thus differing from the criterion (given purely by a 20% rise in PSA) used here. This rate of progression is a result of a mathematical steady state solution to the Lotka–Volterra system (given in the Supplementary Information), where a fully-resistant tumor will remain stable, neither growing nor shrinking under drug treatment. From this we may formulate a mathematical progression limit where the stable tumor size is smaller than threshold for progression, meaning these patients would never reach progression under our criterion, independent of the treatment schedule provided to them. This is observed in 70% of patients, while the remainder have sufficiently slow tumor growth that progression does not occur within clinically feasible timescales, taking over 100 years in some cases. These cases may be interpreted clinically as benign tumors/stable disease, as opposed to the more aggressive, recurrent cases considered in this paper. We therefore only train the DRL framework on the seven virtual patients who are expected to reach progression within 5000 days, since they are most significantly impacted by the limitations of the current treatment paradigms.

Plotting this cohort as a whole in Figure 5a, we observe a broadly negative correlation between cost and turnover; we attribute this to lower turnovers being associated with increasingly competitive sensitive cell populations, thereby raising the relative cost of resistance. Within the whole cohort, these seven patients tend to have a lower total cost and turnover, indicating a more aggressive tumor with a lower expected TTP. We can characterise the tumor in this cost–turnover parameter space (varying tumor dynamics for an single initial tumor profile) more broadly, plotting the expected TTP for each combination of cost and turnover in the first panel of Figure 5c. Again, we see that patients with a lower summed cost and progression have a lower TTP, while all combinations beyond the mathematical progression limit are observed to cycle indefinitely.

**Figure 5:**
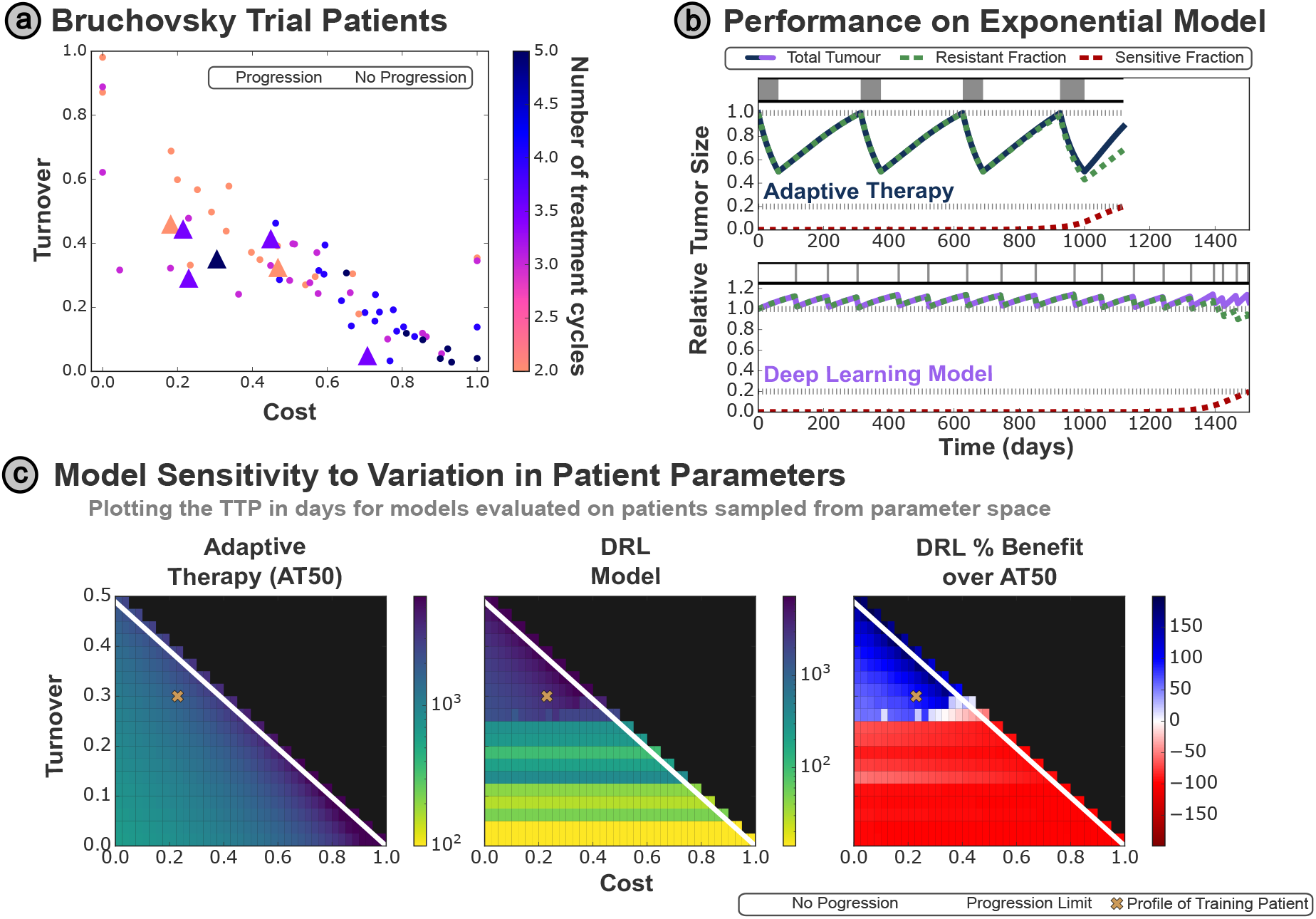
**(a)** Patient parameter values in cost-turnover parameter space from fits conducted by Strobl et al. [17] on patients from the Bruchovsky [47] trial. The patients who progress within 5000 days of AT are denoted by triangles. **(b)** Performance of a pre-trained DRL network when evaluated on a different tumor model (developed by Lu et al. [48]) with modified dynamics (achieved through use of exponential growth terms, and where the progression criterion is based on the resistant subpopulation alone). A DRL framework evaluating treatment decisions weekly is able to consistently outperform AT. **(c)** TTP for a grid of patient profiles under both AT and a DRL framework, which was trained on a single profile (pink cross). The DRL framework is robust to variation in cost, but sensitive to decreases in turnover, where it under-performs relative to AT.

We can now use this continuous space of virtual patients to explore how the DRL framework performs over a range of tumor dynamics that it has not encountered in training. We evaluate the DRL framework on every combination of cost and turnover, comparing a diverse range of virtual patient dynamics using the network trained on the original patient with non-zero cost and turnover (Figure 5c). We see that the DRL is robust to variation in the resistance cost associated with the tumor, being able to deliver equally effective treatment to virtual patients with significantly different resistance costs to the training patient. However, the DRL framework is very sensitive to reductions in the tumor turnover (i.e. the ratio between death and growth rates in the tumor), performing worse than AT50 for patients with slightly reduced turnover (corresponding to faster cell proliferation rates). We attribute this to faster-than-anticipated tumor growth, relative to the growth rate of the training patient, leading to premature progression between treatment decisions. However, the ability of the DRL framework to outperform AT50 for patient profiles with equal or greater turnover than the training patient serves to demonstrate the partial robustness of our framework to uncertainty in patient parameters.

Beyond uncertainty in the tumor’s parameters, we cannot expect all tumors to behave in a way consistent with the virtual patient model (1), given the heterogeneity observed clinically in tumor dynamics and treatment response. We therefore considered the robustness of the DRL framework to variation in the underlying model and therefore tumor dynamics, by evaluating our framework on a modified Lotka–Volterra model introduced by Lu et al. [48] and reproduced in Supplementary Information 5.2. This model uses exponential growth instead of logistic, such that the tumor may grow indefinitely, with a modified progression criterion based on the growth of the resistant subpopulation alone. Despite this significant change in dynamics and progression criterion, we observe that the DRL attains a TTP of 1506 *±* 3 days under weekly treatment evaluation, outperforming the AT50 TTP of 1119 days (95% confidence interval: [1110d, 1128d]).

In the clinic, any treatment framework will be exposed to a range of patients, and should adapt appropriately to their differing characteristics. In this section we have introduced a virtual cohort of patients based on a clinical trial dataset, to test the adaptability of our DRL framework (trained on a single patient parameter set) to a range of treatment responses. We found that the DRL framework could significantly outperform AT50 when evaluated on a different virtual patients with different progression criterion, and broadly outperformed AT50 when the dynamic parameters of the tumor were varied. However, the DRL framework under-performed when treating profiles with an increased tumor growth rate relative to the original training patient.

### 3.5 Personalised treatment though combining mechanistic and DRL frameworks

As we have just shown, a DRL framework that only ‘sees’ one particular patient profile in training has a limited ability to adapt to different patients in evaluation. To expand the framework’s robustness to such variation in the underlying tumor parameters, we will now expose the DRL framework to multiple patient profiles during training. We form a cohort of virtual patients by randomly sampling from the cost-turnover space to represent patients with the same initial tumor but varying underlying dynamics. Importantly, this DRL framework was not trained from scratch; instead we utilize a strategy in machine learning known as ‘transfer learning’, where a pre-existing network is re-trained for a new task. In this case, we started with the single patient network, enabling the DRL framework to develop new strategies based on our sigmoidal treatment strategy (Figure 4a). To capture the uncertainty in patient parameters that we encounter in the clinic, each worker in the DRL network is randomly allocated a new patient for each iteration - the networks are not provided with any further information about that particular patient’s tumor dynamics and so must infer these while providing treatment.

Such uncertainty in patient dynamics can typically be challenging, however our DRL framework was able to match or outperform AT50 for all patients that it had encountered in training. To verify that our framework had not become over-specialised to this new, virtual training cohort, we also evaluated our DRL framework on a set of 20 new virtual patients, drawn from the same region of parameter space. Equivalent performance was also achieved on this set, validating out success is more general and not limited to the specific patient profiles used in training (Figure 6a).

**Figure 6:**
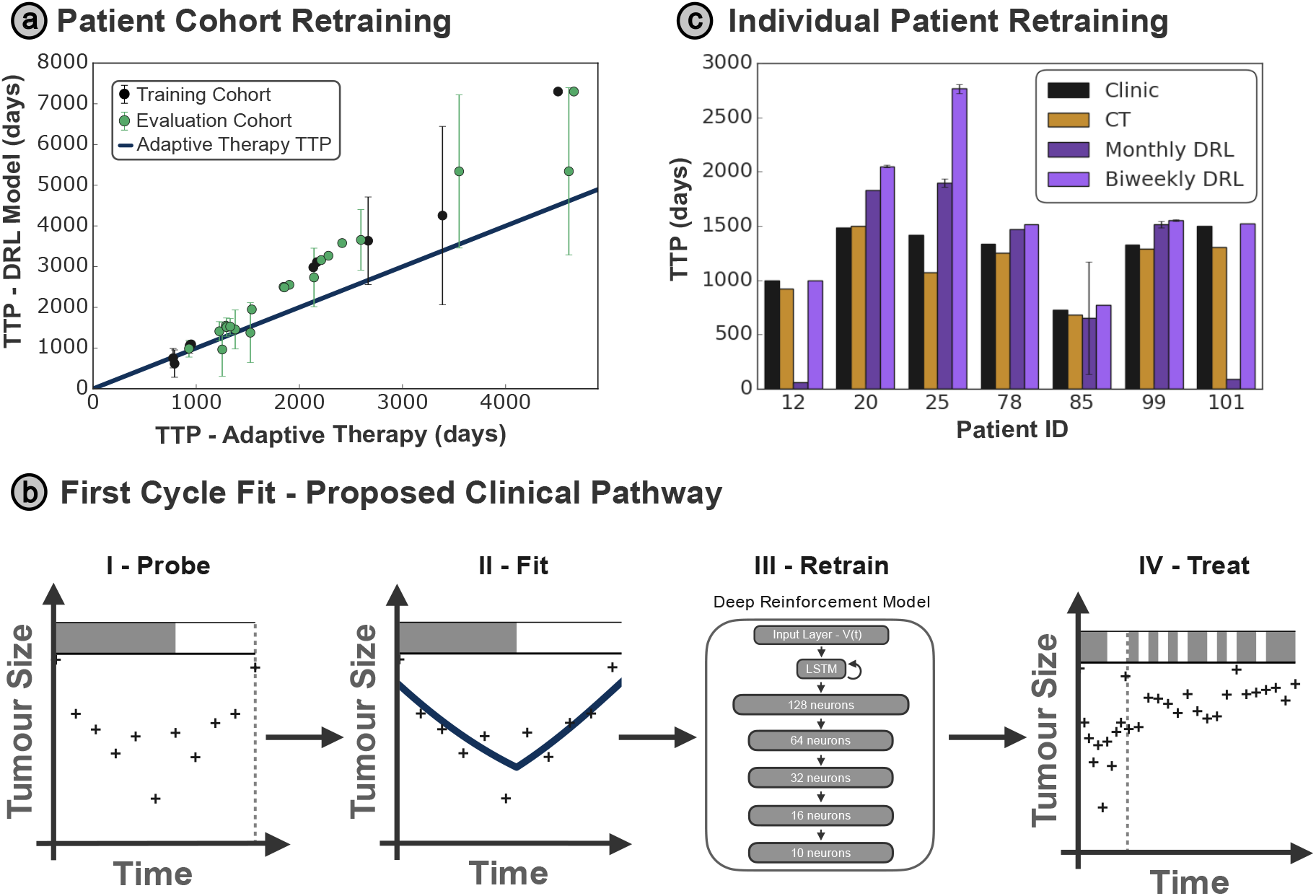
**(a)** Average outcomes of the 10 virtual patients used in training, with variation in each patient’s outcome denoted by error bars corresponding to two standard deviations, compared to 20 further evaluation patients. The DRL framework is able to match or outperform AT for all patients. **(b)** We propose a four-step clinical framework for personalised treatment schedules. Patients undergo an initial ‘probing’ cycle of AT50, to which the virtual patient model is fitted, generating a set of tumor parameters specific to each patient. A copy of the generalised DRL model is then retrained on these personalised parameters, finetuning the network to that patient’s treatment response. The DRL network then provides personalised schedule recommendations throughout the remainder of the treatment schedule. **(c)** Performance of this individualised DRL framework, retraining the generalised DRL network using estimated fits for patients from the Bruchovsky trial. The DRL framework consistently outperforms the standard of care over a biweekly treatment interval, as well as the TTP recorded for the patients in the clinic.

However, such generalist approaches are limited by the variation in virtual patient dynamics, and this may become more significant when considering clinical datasets such as the Bruchovsky cohort, which displays significant variation in patient dynamics (Figure 5a). To improve our generalist framework further, we will explicitly personalise our treatment strategy to each individual, to finetune our network to each patient’s tumor dynamics. In practice, we cannot directly know a patient’s characteristics (such as how fast the tumor will respond to treatment) when they first present in the clinic, preventing us from refining our generalist network on an individual patient profile. One solution we propose here it to approximate each patient’s tumor dynamics by using a ‘probing cycle’ of AT50, throughout which tumor burden metrics are recorded and subsequently fitted to our virtual patient model. This will generate an estimated patient profile upon which we can retrain our DRL framework, to determine future treatment decisions. We present the results of this approach (Figure 5b) as a clinically feasible implementation of our DRL treatment scheduling framework.

While retaining a clinically realistic monthly interval between treatment decisions, we train a collection of DRL frameworks that are able to outperform AT50 for all patients (Figure 6c). Crucially, the retraining of our generalised DRL network is feasible in a clinical setting, achieving sufficient personalisation to each patient after as little as 2000 epochs, taking approximately 20 minutes to train on a standard laptop (Intel i5, 4 cores, 1.70GHz). This approach heralds a new avenue in personalised medicine, through tailored treatment schedules on a per-patient basis, driven by their initial response to one cycle of treatment.

## 4 Discussion

Following the recent clinical implementation of adaptive therapy on castrate-resistant prostate cancer by Zhang et al. [23], and multiple ongoing adaptive therapy trials in skin (NCT05651828), prostate (NCT05393791) and ovarian (NCT05080556) cancers, it is of significant interest to better understand if we can optimise adaptive treatment strategies above and beyond the classic AT50 approach. While such optimisation may be conducted analytically where binary treatment decisions are applied to simple models, they quickly become infeasible in more complex cases. We therefore consider the application of deep learning models, introducing a reinforcement learning framework wherein the DRL framework interacts with a virtual patient model to develop successful treatment strategies based only on a single clinically accessible tumor metric: total burden.

We demonstrate this DRL framework can outperform the ‘rule-of-thumb’ adaptive strategy (AT50) used clinically by Zhang et al. [23]. This strategy is robust to varying model parameterisations, and across multiple underlying tumor models. We uncovered a novel relationship in regards to the frequency of treatment decisions (i.e. how often patient data are collected and the current treatment reevaluated), showing that application of treatment should occur at lower frequencies in initial training, to allow the DRL framework to learn meaningful and interpretable treatment strategies. However, this must be balanced by sufficient time to react to tumour growth, with faster growing tumors requiring more frequent treatment decisions.

Exploring the decision making process behind the DRL network, we discovered that it had learnt to mimic optimal strategies previously derived through optimal control theory [7, 10, 29], maintaining a large sensitive cell population for maximal suppression of the resistant cells while avoiding the progression limit on tumor size. This trade-off discovered by the DRL framework motivated an analytic exploration of the optimal treatment strategy, which in turn produced a functional relationship between tumor size and treatment timing that accurately matched the treatment threshold learnt by the DRL framework. Given this framework had no underlying knowledge of the mathematical representation of the system, this illustrates the power of DRL approaches to study more complex models or treatment paradigms (such as multiple treatment drugs or non-binary dosing levels), where mathematical analysis may not be feasible.

The dependence of this optimal treatment strategy on individual patient parameters, alongside the significant variation in the adaptive cycling dynamics observed between patients in the clinic (Figure 1a), highlights the importance of decision-making frameworks that are robust to variation between patients and uncertainty in individual patient dynamics. By retraining the DRL network on a cohort of 10 virtual patient profiles, it was able to match or outperform AT50 in 95% of cases, from a cohort of testing patients it had never encountered previously. Finally, we propose a pathway to integrate mechanistic modelling with DRL to tailor such robust, generalist DRL strategies to individual patients. Using a probing cycle to characterise the treatment response of each patient within our virtual patient model, we are able to retrain the DRL network on each patient’s dynamics (with limited computational resources and expertise in the clinic), to produce a personalised treatment schedule that consistently outperforms clinical standard-of-care protocols as well as AT50.

In future work, we plan to further leverage uncertainty about patient measurements and dynamics by generating a virtual cohort of patients for each individual patient, then we will use this cohort to retrain our more generalized DRL framework for that individual patient. These virtual cohorts would match the real patient’s dynamics within a given error similar to the phase i approach we have previously used [49, 50]. Such approaches would retain the robustness benefits from training on a range of patients, while allowing treatment schedules to be tailored to individual patients based on their profiles. Practically, this would require minimal retraining of a generalist model using a best estimate of the tumor parameters from fitting to the patient’s current clinical history. The objective function also allows for an alternative route to personalise treatment; by modifying the reward to punish cumulative drug dosage over a given level, treatment schedules could be adapted for patients struggling to complete their planned course due to side-effects/cytotoxicity.

The approach we have presented here is not the first DRL framework to tackle adaptive therapy. The work of Lu et al. [48] is focused solely on prostate cancer and used a different underlying model, but importantly trained the network on both the PSA as well as the senstitve and resistant populations; while PSA is easily measured in a blood draw there is no direct way to measure the numbers of sensitive and resistant cells in a real patient. While they were also able to produce significantly improved responses for a single patient, the generalisability of our approach, and the clear path for translation, make our implementation somewhat more robust. In addition, our novel discovery of the analytical relationship between tumor size and treatment timing for the treatment threshold sets our work apart in terms of interpretability, but also in regards to the frequency of treatments. With the ongoing revolution in wearables, a future where tumor burden could be measured in real time isn’t so unrealistic. Our results imply one must be cautious in making too frequent treatment decisions in DRL training, since this effectively reduces the impact of each decision and may increase the impact of noise on the system.

The approaches presented here are highly generalisable to other forms of cancer, and applicable in any context where total tumor burden or drug load must be managed, either to prevent treatment resistance or to reduce adverse side-effects. They may also be extended to multi-drug paradigms, or to allow variable dosing levels for individual drugs, provided a dose response function is known. The application of DRL frameworks therefore allows for optimisation in a wide range of clinical settings where clinical standards are currently determined by a ‘best guess’ that, in many cases, may be far from the optimal strategies a DRL framework could obtain.

## References

[1] K. Bukowski, M. Kciuk, and R. Kontek, “Mechanisms of multidrug resistance in cancer chemotherapy,” International Journal of Molecular Sciences, vol. 21, p. 3233, May 2020.

[2] C. Holohan, S. V. Schaeybroeck, D. B. Longley, and P. G. Johnston, “Cancer drug resistance: an evolving paradigm,” Nature Reviews Cancer, vol. 13, pp. 714–726, Sept. 2013.

[3] K. Ganesh and J. Massagué, “Targeting metastatic cancer,” Nature Medicine, vol. 27, pp. 34–44, Jan. 2021.

[4] A. J. Tevaarwerk, R. J. Gray, B. P. Schneider, M. L. Smith, L. I. Wagner, J. H. Fetting, N. Davidson, L. J. Goldstein, K. D. Miller, and J. A. Sparano, “Survival in patients with metastatic recurrent breast cancer after adjuvant chemotherapy: little evidence of improvement over the past 30 years,” Cancer, vol. 119, no. 6, pp. 1140–1148, 2013.

[5] M. C. Perry, The Chemotherapy Source Book, vol. 117. Lippincott Williams & Wilkins, 1992.

[6] C. C. Maley, A. Aktipis, T. A. Graham, A. Sottoriva, A. M. Boddy, M. Janiszewska, A. S. Silva, M. Gerlinger, Y. Yuan, K. J. Pienta, K. S. Anderson, R. Gatenby, C. Swanton, D. Posada, C. I. Wu, J. D. Schiffman, E. S. Hwang, K. Polyak, A. R. Anderson, J. S. Brown, M. Greaves, and D. Shibata, “Classifying the evolutionary and ecological features of neoplasms,” Nature Reviews Cancer, vol. 17, pp. 605–619, 2017.

[7] R. Martin, M. Fisher, R. Minchin, and K. Teo, “Optimal control of tumor size used to maximize survival time when cells are resistant to chemotherapy,” Mathematical Biosciences, vol. 110, pp. 201–219, July 1992.

[8] H. C. Monro and E. A. Gaffney, “Modelling chemotherapy resistance in palliation and failed cure,” Journal of Theoretical Biology, vol. 257, pp. 292–302, Mar. 2009.

[9] R. A. Gatenby, A. S. Silva, R. J. Gillies, and B. R. Frieden, “Adaptive therapy,” Cancer Research, vol. 69, pp. 4894–4903, June 2009.

[10] E. Hansen, R. J. Woods, and A. F. Read, “How to use a chemotherapeutic agent when resistance to it threatens the patient,” PLOS Biology, vol. 15, p. e2001110, Feb. 2017.

[11] R. A. Gatenby, “A change of strategy in the war on cancer,” Nature, vol. 459, pp. 508–509, May 2009.

[12] R. A. Gatenby and J. S. Brown, “The evolution and ecology of resistance in cancer therapy,” Cold Spring Harbor Perspectives in Medicine, vol. 10, p. a040972, Nov. 2020.

[13] R. J. Gillies, D. Verduzco, and R. A. Gatenby, “Evolutionary dynamics of carcinogenesis and why targeted therapy does not work,” Nature Reviews Cancer, vol. 12, pp. 487–493, June 2012.

[14] J. A. Gallaher, P. M. Enriquez-Navas, K. A. Luddy, R. A. Gatenby, and A. R. Anderson, “Spatial heterogeneity and evolutionary dynamics modulate time to recurrence in continuous and adaptive cancer therapies,” Cancer Research, vol. 78, pp. 2127–2139, Jan. 2018.

[15] K. Bacevic, R. Noble, A. Soffar, O. W. Ammar, B. Boszonyik, S. Prieto, C. Vincent, M. E. Hochberg, L. Krasinska, and D. Fisher, “Spatial competition constrains resistance to targeted cancer therapy,” Nature Communications, vol. 8, Dec. 2017.

[16] M. A. R. Strobl, J. Gallaher, J. West, M. Robertson-Tessi, P. K. Maini, and A. R. A. Anderson, “Spatial structure impacts adaptive therapy by shaping intra-tumoral competition,” Communications Medicine, vol. 2, pp. 1–18, 4 2022.

[17] M. A. Strobl, J. West, Y. Viossat, M. Damaghi, M. Robertson-Tessi, J. S. Brown, R. A. Gatenby, P. K. Maini, and A. R. Anderson, “Turnover modulates the need for a cost of resistance in adaptive therapy,” Cancer Research, vol. 81, pp. 1135–1147, Feb. 2021.

[18] N. Farrokhian, J. Maltas, M. Dinh, A. Durmaz, P. Ellsworth, M. Hitomi, E. McClure, A. Marusyk, A. Kaznatcheev, and J. G. Scott, “Measuring competitive exclusion in non-small cell lung cancer,” Science Advances, vol. 8, 2022.

[19] P. M. Enriquez-Navas, Y. Kam, T. Das, S. Hassan, A. Silva, P. Foroutan, E. Ruiz, G. Martinez, S. Minton, R. J. Gillies, and R. A. Gatenby, “Exploiting evolutionary principles to prolong tumor control in preclinical models of breast cancer,” Science Translational Medicine, vol. 8, Feb. 2016.

[20] J. Wang, Y. Zhang, X. Liu, and H. Liu, “Optimizing adaptive therapy based on the reachability to tumor resistant subpopulation,” Cancers, vol. 13, 2021.

[21] I. Smalley, E. Kim, J. Li, P. Spence, C. J. Wyatt, Z. Eroglu, V. K. Sondak, J. L. Messina, N. A. Babacan, S. S. Maria-Engler, L. D. Armas, S. L. Williams, R. A. Gatenby, Y. A. Chen, A. R. Anderson, and K. S. Smalley, “Leveraging transcriptional dynamics to improve braf inhibitor responses in melanoma,” EBioMedicine, vol. 48, pp. 178–190, 2019.

[22] J. Zhang, J. J. Cunningham, J. S. Brown, and R. A. Gatenby, “Integrating evolutionary dynamics into treatment of metastatic castrate-resistant prostate cancer,” Nature Communications, vol. 8, Nov. 2017.

[23] J. Zhang, J. Cunningham, J. Brown, and R. Gatenby, “Evolution-based mathematical models significantly prolong response to abiraterone in metastatic castrate-resistant prostate cancer and identify strategies to further improve outcomes,” eLife, vol. 11, June 2022.

[24] M. Ferńandez-Cancio, N. Camats, C. Flück, A. Zalewski, B. Dick, B. Frey, R. Monné, N. Torán, L. Audi, and A. Pandey, “Mechanism of the dual activities of human CYP17A1 and binding to anti-prostate cancer drug abiraterone revealed by a novel V366M mutation causing 17, 20 lyase deficiency,” Pharmaceuticals, vol. 11, p. 37, Apr. 2018.

[25] P. Therasse, S. G. Arbuck, E. A. Eisenhauer, J. Wanders, R. S. Kaplan, L. Rubinstein, J. Verweij, M. V. Glabbeke, A. T. van Oosterom, M. C. Christian, and S. G. Gwyther, “New guidelines to evaluate the response to treatment in solid tumors,” JNCI: Journal of the National Cancer Institute, vol. 92, pp. 205–216, Feb. 2000.

[26] F. H. Schröder, I. van der Cruijsen-Koeter, H. J. de Koning, A. n. Vis, R. F. Hoedemaeker, and R. Kranse, “Prostate cancer detection at low prostate specific antigen,” Journal of Urology, vol. 163, pp. 806–812, Mar. 2000.

[27] R. Lieberman, “Evidence-based medical perspectives: The evolving role of PSA for early detection, monitoring of treatment response, and as a surrogate end point of efficacy for interventions in men with different clinical risk states for the prevention and progression of prostate cancer,” American Journal of Therapeutics, vol. 11, pp. 501–506, Nov. 2004.

[28] E. Hansen and A. F. Read, “Modifying adaptive therapy to enhance competitive suppression,” Cancers, vol. 12, pp. 1–13, 2020.

[29] Y. Viossat and R. Noble, “A theoretical analysis of tumour containment,” Nature Ecology & Evolution, vol. 5, pp. 826–835, Apr. 2021.

[30] E. Kim, J. S. Brown, Z. Eroglu, and A. R. Anderson, “Adaptive therapy for metastatic melanoma: Predictions from patient calibrated mathematical models,” Cancers, vol. 13, p. 823, Feb. 2021.

[31] R. Brady-Nicholls and H. Enderling, “Range-bounded adaptive therapy in metastatic prostate cancer,” Cancers 2022, Vol. 14, Page 5319, vol. 14, p. 5319, 10 2022.

[32] Y. LeCun, Y. Bengio, and G. Hinton, “Deep learning,” Nature, vol. 521, pp. 436–444, May 2015.

[33] K. Munir, H. Elahi, A. Ayub, F. Frezza, and A. Rizzi, “Cancer diagnosis using deep learning: A bibliographic review,” Cancers, vol. 11, p. 1235, Aug. 2019.

[34] D. Silver, A. Huang, C. J. Maddison, A. Guez, L. Sifre, G. van den Driessche, J. Schrittwieser, I. Antonoglou, V. Panneershelvam, M. Lanctot, S. Dieleman, D. Grewe, J. Nham, N. Kalchbrenner, I. Sutskever, T. Lillicrap, M. Leach, K. Kavukcuoglu, T. Graepel, and D. Hassabis, “Mastering the game of go with deep neural networks and tree search,” Nature, vol. 529, pp. 484–489, Jan. 2016.

[35] M. R. Bachute and J. M. Subhedar, “Autonomous driving architectures: Insights of machine learning and deep learning algorithms,” Machine Learning with Applications, vol. 6, p. 100164, Dec. 2021.

[36] A. Esteva, A. Robicquet, B. Ramsundar, V. Kuleshov, M. DePristo, K. Chou, C. Cui, G. Corrado, S. Thrun, and J. Dean, “A guide to deep learning in healthcare,” Nature Medicine, vol. 25, pp. 24–29, Jan. 2019.

[37] Y. Liu, B. Logan, N. Liu, Z. Xu, J. Tang, and Y. Wang, “Deep reinforcement learning for dynamic treatment regimes on medical registry data,” in 2017 IEEE International Conference on Healthcare Informatics (ICHI), IEEE, Aug. 2017.

[38] D. Engelhardt, “Dynamic control of stochastic evolution: A deep reinforcement learning approach to adaptively targeting emergent drug resistance,” Journal of Machine Learning Research, vol. 21, no. 203, pp. 1–30, 2020.

[39] D. T. Weaver, J. Maltas, and J. G. Scott, “Reinforcement learning informs optimal treatment strategies to limit antibiotic resistance,” bioRxiv, Jan. 2023.

[40] J.-N. Eckardt, K. Wendt, M. Bornhäuser, and J. M. Middeke, “Reinforcement learning for precision oncology,” Cancers, vol. 13, p. 4624, Sept. 2021.

[41] C. Yu, J. Liu, S. Nemati, and G. Yin, “Reinforcement learning in healthcare: A survey,” ACM Computing Surveys, vol. 55, pp. 1–36, Jan. 2023.

[42] Y. Zhao, M. R. Kosorok, and D. Zeng, “Reinforcement learning design for cancer clinical trials,” Statistics in Medicine, vol. 28, pp. 3294–3315, Nov. 2009.

[43] C. Maier, N. Hartung, C. Kloft, W. Huisinga, and J. Wiljes, “Reinforcement learning and bayesian data assimilation for model-informed precision dosing in oncology,” CPT: Pharmacometrics and Systems Pharmacology, vol. 10, pp. 241–254, Mar. 2021.

[44] B. Eastman, M. Przedborski, and M. Kohandel, “Reinforcement learning derived chemotherapeutic schedules for robust patient-specific therapy,” Scientific Reports, vol. 11, Sept. 2021.

[45] V. Mnih, A. P. Badia, M. Mirza, A. Graves, T. Lillicrap, T. Harley, D. Silver, and K. Kavukcuoglu, “Asynchronous methods for deep reinforcement learning,” in Proceedings of The 33rd International Conference on Machine Learning (M. F. Balcan and K. Q. Weinberger, eds.), vol. 48 of Proceedings of Machine Learning Research, (New York, New York, USA), pp. 1928–1937, PMLR, June 2016.

[46] Y. Gao, Y. Liu, H. Zhang, Z. Li, Y. Zhu, H. Lin, and M. Yang, Estimating GPU Memory Consumption of Deep Learning Models, p. 1342–1352. New York, NY, USA: Association for Computing Machinery, 2020.

[47] N. Bruchovsky, L. Klotz, J. Crook, S. Malone, C. Ludgate, W. J. Morris, M. E. Gleave, and S. L. Goldenberg, “Final results of the canadian prospective phase II trial of intermittent androgen suppression for men in biochemical recurrence after radiotherapy for locally advanced prostate cancer,” Cancer, vol. 107, no. 2, pp. 389–395, 2006.

[48] Y. Lu, Q. Chu, Z. Li, M. Wang, and Q. Zhang, “Deep reinforcement learning identifies personalized intermittent androgen deprivation therapy for prostate cancer,” May 2022.

[49] E. Kim, V. W. Rebecca, K. S. Smalley, and A. R. Anderson, “Phase i trials in melanoma: A framework to translate preclinical findings to the clinic,” European Journal of Cancer, vol. 67, pp. 213–222, Nov. 2016.

[50] M. Robertson-Tessi, J. S. Brown, M. I. Poole, M. Johnson, A. Marusyk, J. A. Gallaher, K. A. Luddy, C. J. Whelan, J. West, M. Strobl, V. Turati, H. Enderling, M. J. Schell, A. Tan, T. Boyle, R. Makanji, J. Farinhas, H. Soliman, D. Lemanne, R. A. Gatenby, D. R. Reed, A. R. A. Anderson, and C. H. Chung, “Feasibility of an evolutionary tumor board for generating novel personalized therapeutic strategies,” Jan. 2023.

