## Supplementary Information for "Learning to Adapt - Deep Reinforcement Learning in Treatment-Resistant Prostate Cancer"

Jan 2023

#### 1 Deep Learning Methods

This section provides further exposition and psuedocode implementations to supplement the explanation of the DRL model in Section 2.3.

We transform cancer treatment into a reinforcement learning problem, in which a computational agent learns to make decisions in an environment by trial and error based on some reward function, allowing agents to learn from unstructured input data [1]. We construct this as a 'model-free' problem, where information about the dynamics of the environment is not given explicitly, but instead inferred through interactions with the environment. We adopt an Actor-Critic method, which combines the advantages of policy and value based methods, whereby an 'Actor' updates the policy distribution according to a value function estimated by the 'Critic' [2]. An illustration of this process is given in pseudocode in Algorithm 1.

**Algorithm 1** Actor–Critic Method

**Require:** Parameters: $\alpha$ (learning rate), $\gamma$ (discount factor)
**Require:** Initial values: state $s$, policy parameters $\theta$, value $w$ and action $a$.

**for** $t \leftarrow 1...T$ **do**
    Calculate reward $r$ from reward function $R(s, a)$
3:    Compute next state $s'$ based on previous $P(s'|s, a)$
    Sample next action $a'$ according to policy $\pi_\theta(a'|s')$
    $\theta \leftarrow \theta + \alpha Q_w(s, a)\nabla_\theta \log \pi_\theta(a|s)$ ▷ Update policy parameters[1]
6:    $\delta = r + \gamma Q_w(s', a') - Q_w(s, a)$ ▷ Compute correction for action-value
    $w \leftarrow w + \alpha\delta\nabla_w Q_w(s, a)$ ▷ Update parameters $w$ of value function $Q_w$
    $a \leftarrow a'$; $s \leftarrow s'$
9: **end for**

Note that Operation 5 utilises a baseline comparison for the value of the cumulative reward; this results in a smaller absolute value, which reduces the error in gradient-based updates. This choice of baseline is taken from Q Actor-Critic [4], but it is not the only option; a popular alternative is advantage Actor-Critic [5]. Here the baseline $A_w(s, a)\nabla_\theta \log \pi_\theta(a|s)$ is used, where the advantage value $A_w(s, a)$ is given by:

$$A_w(s, a) = Q_w(s, a) - V(s), \tag{1}$$

i.e the added benefit from taking the given action $a$ from state $s$ compared to the expected value $V$ (based on all actions) from state $s$.

As with all GPU-based deep-learning algorithms, this typically has a high computational cost, and requires specialist architecture to train performant models. We therefore utilise the

[1] This uses the standard form for the policy gradient [3].

asynchronous, advantage Actor–Critic (A3C) framework pioneered by Mnih et al. [6] which combines the Advantage Actor-Critic method with a lightweight CPU framework supporting parallel actor-learning training asynchronously. The policy and value functions are not updated every timestep, and share a convolutional neural network framework with separate softmax and linear outputs for the policy and value, respectively. Since the threads are asynchronous, they update from the master policy at different times, and so have slightly different policies exploring different regions of the environment. A simplistic pseudocode representation of each actor-learner thread is given in Algorithm 2, adapted from [6].

**Algorithm 2** Asynchronous Advantage Actor–Critic - adapted from [6]

**Require:** Parameters: $t_{max}$ (update rate), $T_{max}$ (iteration number), $\gamma$ (discount factor)
**Require:** Global shared variables: $\theta, \theta_v$ (parameter vectors), $T$ (counter)
**Require:** Thread-specific variables: $\theta, \theta_v$ (parameter vectors)

```
t ← 1                                                      ▷ Initialise thread step counter
while T < T_max do
    Reset gradients dθ ← 0; dθ_v ← 0
    Synchronise with global parameters θ' ← θ; θ'_v ← θ_v
    t_start = t
    Obtain state s_t
    while t − t_start < t_max do
        Perform action a_t according to policy π(a_t|s_t; θ')
        Receive reward r_t and new state s_{t+1}
        t ← t + 1
        T ← T + 1
    end while
    R = V(s_t, θ'_v)                                       ▷ Bootstrap from last state
    for i ∈ {t − 1, ..., t_start} do
        R ← r_i + γR
        dθ ← dθ + ∇_θ' log π(a_i|s_i; θ')(R − V(s_i; θ'_v))     ▷ Accumulate gradients wrt θ'
        dθ_v ← dθ_v + ∂(R − V(s_i; θ'_v))^2/∂θ'_v
    end for
    Asynchronous update of θ using dθ and of θ_v using dθ_v
end while
```

Within this, each worker network is constructed as follows:

1. Input Layers
    - Long Short-Term Memory layer [7] gives 4-dimensional output.
2. Hidden Layers
    - Fully connected layers for each of the sizes: [128, 64, 32, 16, 10].
    - Each layer is multiplied by the previous output to produce a tensor of hidden units.
    - Use a rectified linear activation function.
3. Output Layers
    - (a) Policy - Fully connected layer of output size 2, softmax activation function [8].
    - (b) Value - Fully connected layer of output size 1, linear activation function.

Note that the output size of the policy is determined by the number of policy options available; this assumes a binary treatment decision (i.e. treatment is either given or withheld).

#### 2 Virtual Patient Parameters

The parameters used in the virtual patient model and taken from Strobl et al. [9] and outlined in Table 1.

| Name | Description | Value/Range | Reference |
|---|---|---|---|
| $r_S$ | Sensitive cell proliferation rate | $0.027\,day^{-1}$ | Adopted from [10] |
| $r_R$ | Resistant cell proliferation rate | $0.5r_S$ - $1.0r_S$ | Lower limit [11], upper limit of no cost |
| $d_S, d_R$ | Natural cell death rate | $0.0r_S$ - $0.5r_S$ | Lower limit given by zero turnover<br>Upper limit adopted from [12] |
| $d_D$ | Drug-induced cell killing | 1.5 | Adopted from [13] |
| $N_0$ | Initial tumor cell density | $0.1 - 0.75$ | Values in this range reported by [14] |
| $R_0$ | Initial resistant cell fraction | $0.001N_0 - 0.1N_0$ | Values in this range reported by [15] |

Table 1: Parameter values/ranges used for the virtual patient.

#### 3 Progression Analysis

##### 3.1 Progression with Treatment

As discussed in Section 3.4, the Lotka–Volterra model (Section 2.1 - (1)) allows for a steady state which prevents progression in some cases. We will first derive this under continuous (MTD) therapy, where the sensitive population will quickly be depleted to extinction, leaving a fully resistant population at the steady state. However this final resistant population has no dependence on the drug concentration, and so this steady state may be ultimately achieved under any treatment strategy.

Our original model for the virtual patient is reproduced below:

$$\begin{aligned}\frac{dS}{dt} &= r_S S\left(1-\frac{S+R}{K}\right)\times(1-d_D D)-dS\\ \frac{dR}{dt} &= r_R R\left(1-\frac{S+R}{K}\right)-dR\end{aligned} \tag{1 revisited}$$

When $S(t)=0$ this reduces to:

$$\frac{dR}{dt} = r_R R\left(1-\frac{R}{K}\right)-dR, \tag{2}$$

which in steady state ($\frac{dR}{dt}=0$) defines the steady state resistant cell population $R'$ according to the expression:

$$dR' = r_R R'\left(1-\frac{R'}{K}\right). \tag{3}$$

Simple rearrangement gives the result for $R'$:

$$R' = K\left(1-\frac{d_R}{r_R}\right), \tag{4}$$

i.e. the resistant population reaches a non-zero, constant steady state provided that $r_R > d_r$. This will only result in progression when:

$$1.2(S_0+R_0) < K\left(1-\frac{d_R}{r_R}\right), \tag{5}$$

where $S_0$ and $R_0$ are the initial populations of sensitive and resistant cells respectively.

Patient profiles which obey this condition will therefore be stable indefinitely under a MTD strategy. Any patient who does not progress under MTD will also not progress under the 'rule of thumb' AT strategy, and will either cycle indefinitely, or ultimately experience extinction of the sensitive population to revert back to the fully resistant steady state derived above for MTD.

##### 3.2 Progression without Treatment

It is important to note that these stable profiles are only stable under continuous treatment, and may still undergo progression under sub-optimal strategies. However this can only occur through insufficient treatment of the sensitive population, as the resistant population cannot result in progression alone. In this case, under the assumption that $S_0 >> R_0$, we may neglect the resistant population to derive an equivalent condition to (5). When no treatment is given, patients will only reach progression if:

$$1.2S_0 < K\left(1 - \frac{d_S}{r_S}\right). \tag{6}$$

Otherwise, tumors will maintain a non-zero steady state below the progression threshold, provided $r_S > d_S$ (else the tumor will be eliminated). Note that our formulation of resistance cost requires that $r_S \geq r_R$, while we assume $d_S = d_R$ throughout. This also means that (6) is inherently stricter than (5), i.e. patients who do not progress under continuous treatment may still progress without treatment. It is worth noting that neither requirement depends on the initial resistant fraction, but only compares the initial tumor size to its growth and death rates.

##### 3.3 Cost-Turnover Space

We may reframe condition (5) for progression under treatment in terms of cost-turnover parameter space. The cost is characterised by the relative proliferation rates for sensitive and resistant cells $(1 - r_R/r_S)$, while the cell turnover represents the natural death rate of cells $d/r_s$. Rewriting these in terms of the resistant tumor properties:

$$r_R = r_S(1 - cost); \qquad d_r = r_s \times turnover, \tag{7}$$

we may write (5) as:

$$1.2(S_0 + R_0) < K\left(1 - \frac{turnover}{1 - cost}\right). \tag{8}$$

In cost-turnover space, we only observe progression if:

$$turnover < \left(1 - 1.2\frac{S_0 + R_0}{K}\right)(1 - cost). \tag{9}$$

This line is plotted for reference in Figure 5)c. Naturally, progression cannot occur for $cost = 1$, as this would correspond to a zero proliferation rate for the resistant cells. More interestingly, it is possible to avoid progression even without a resistance cost, provided the carrying capacity is sufficiently small to restrict the logistic growth of the system.

#### 4 Critical Treatment Threshold

Due to the simplicity of this Lotka–Volterra model (Section 2.1 - (1)), we may derive an optimal treatment strategy analytically. This is based on the principle that the growth of the resistant sub-population may only be suppressed by the sensitive sup-population, and so maximising

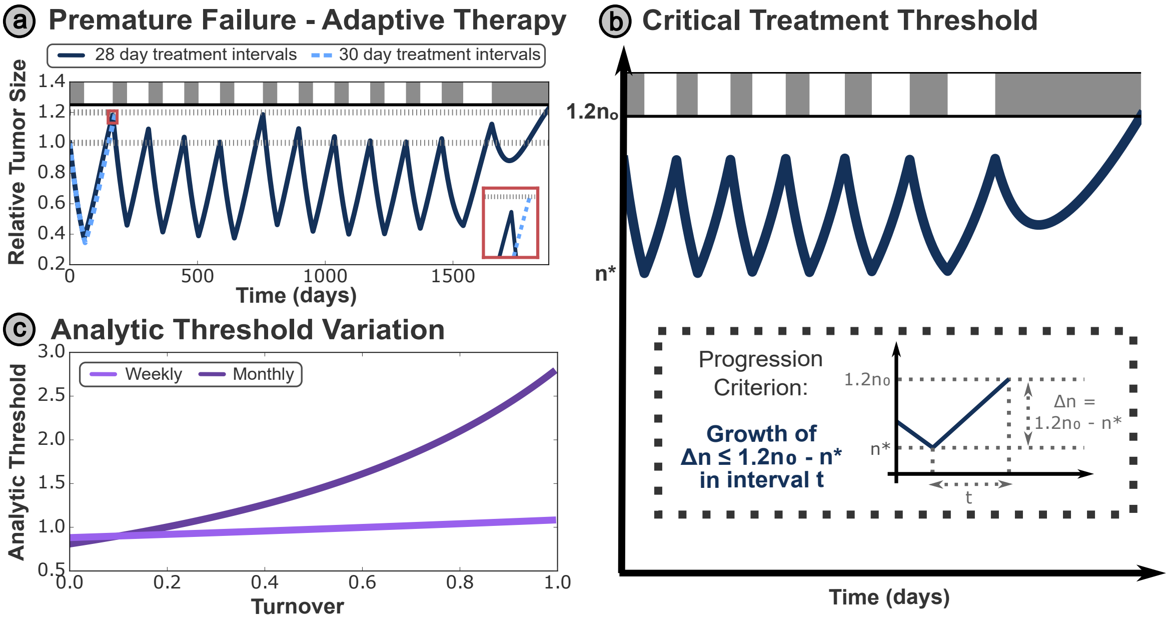


Figure 1: **(a)** Premature failure, where the tumor grows from $n(t) < 0.5n_0$ to $n(t) > 1.2n_0$ in a single treatment cycle may be induced from small increases in the treatment interval. **(b)** We define the time taken by a tumor to grow between the thresholds as the critical treatment interval (as treatment intervals greater than this will result in premature failure). **(c)** The critical treatment threshold varies both with the treatment interval used, and patient parameters such as the turnover.

the sensitive population maximally delays progression due to the uncontrolled growth of the resistant sub-population. This is constrained by the requirement that the sensitive population is controlled such that the total tumor size remains under the progression criterion at all times.

#### 4.1 Optimal Treatment Interval

We will determine the maximal possible treatment interval that prevents premature failure under conventional adaptive therapy. This is directly determined by the growth characteristics of the tumor - specifically failure occurs when the tumor can grow from the treatment threshold size to the progression threshold size within one treatment interval.

These characteristics are highly sensitive to the value of the treatment interval - Figure 1a demonstrates that even small increases in the treatment interval may transform a successful strategy into one that undergoes progression in the first treatment cycle, with the tumor size $n(t)$ jumping from $n(t) < 0.5n_0$ (the threshold for treatment in conventional AT50) to $n(t) > 1.2n_0$ (the threshold for progression) in a single treatment cycle. We will refer to this as premature failure, as the tumor progresses under a treatment holiday due to the growth of the sensitive subpopulation, and this progression would have been prevented by earlier drug treatment. The critical treatment time ($t$ in Figure 1b) is the time for the tumor to grow from one threshold to the other.

We may estimate this critical interval, assuming $R = 0$ such that the total tumor size $n$ is equal to the sensitive population $S$, although we will address this assumption in Section 4.2.

Rewriting (1) with $S \approx n$ while $R \approx 0$, we obtain:

$$\frac{dn}{dt} = r_S n \left(1 - \frac{n}{K}\right) - d_S n. \tag{10}$$

This differential equation is separable:

$$\int_{n^*}^{1.2n_0} \frac{1}{r_S n\left(1-\frac{n}{K}\right) - d_S n} dn = \int_0^{\tau} t = \tau, \tag{11}$$

where we integrate over the critical treatment period. The integral may be separated in partial fractions:

$$\tau = \int_{n^*}^{1.2n_0} \frac{r}{(d_S - r_S)(-Kr_S + r_S n + Kd_S)} + \frac{1}{n(r_S - d_S)} dn, \tag{12}$$

which may be evaluated to obtain:

$$\tau = \frac{1}{r-d}\, ln \left[ \frac{K(r_S - d_S) - r_S n^*}{K(r_S - d_S) - 1.2 r_S n_0} \frac{1.2n_0}{n^*} \right]. \tag{13}$$

##### 4.2 Non-negligible resistant fractions

In the prior sections, we have assumed the resistant cell population ($R$) is negligible; while this may be true initially it cannot hold throughout the entire treatment schedule (as this subpopulation will ultimately grow sufficiently to result in relapse). While such relapse is inevitable in most parameterisations, we must ensure that any late-stage failure is not a product of insufficient treatment. Equivalently, we must show that our critical treatment interval $\tau$ still holds for any tumor composition $n = S + R$.

Trivially for $r_S = rR$; $d_S = d_R$:

$$\frac{dn}{dt} = \frac{dS}{dt} + \frac{dR}{dt} = r_S n \left(1 - \frac{n}{K}\right) - d_S n, \tag{14}$$

which is equivalent to our previous case (as sensitive and resistant cells are behave identically when $D = 0$), and so the result from Section 4.1.

More generally, our model considers tumors with a non-zero resistant cost (i.e. where $r_R < r_S$):

$$\frac{dn}{dt} = (r_S S + r_R R)\left(1 - \frac{n}{K}\right) - d_S n. \tag{15}$$

This equation requires knowledge of the initial tumor composition $S(t = 0), R(t = 0)$, as well as the initial tumor size $n_0 = S(t = 0) + R(t = 0)$. However, as $r_R < r_S$ (a positive resistance cost), $\frac{dn}{dt}$ is maximised in the case where $n_0 = S(t = 0)$, as considered previously. Therefore, the expression (13) still holds for an arbitrary initial tumor composition.

##### 4.3 Optimal Treatment Threshold

In practice, the treatment interval is often determined by clinical availability and practical restrictions. We may however rearrange (13) to obtain an expression for the optimal treatment threshold based on a given treatment interval $\tau$:

$$n^* = \frac{K(d_S - r_S)}{\left(\frac{K(d_S - r_S)}{1.2n_0} + r\right) e^{(r_S - d_S)\tau} - r_S} \tag{16}$$

This threshold increases with the critical treatment interval $\tau$, and also with the cell turnover in the tumor (Figure 1c).

#### 5 Further DRL Results

##### 5.1 DRL Variability

In Section 3.1, we present results obtained by training the DRL model on a single patient. These are averaged over 100 evaluations, to account for the stochasticity in patient outcomes. This stochasticity is not a product of the virtual patient itself (whose treatment response is calculated by a deterministic set of differential equations), but rather inherent in the decision making process of the DRL framework, such that each iteration receives a slightly different treatment schedule.

We characterise this stochasticity in Figure 2, showing significant variation between individual evaluations of the patient, with the shortest TTP almost a third of the longest, for the same patient profile. This variation means that this patient would have a significant probability of performing worse on the DRL strategy compared to AT50, despite the DRL framework having a mean TPP over 200 days greater. In Section 3.2, we subsequently demonstrate how variation in the performance of the DRL model can be reduced by increasing the interval between treatment decisions, enabling the DRL framework to consistently outperform AT50.

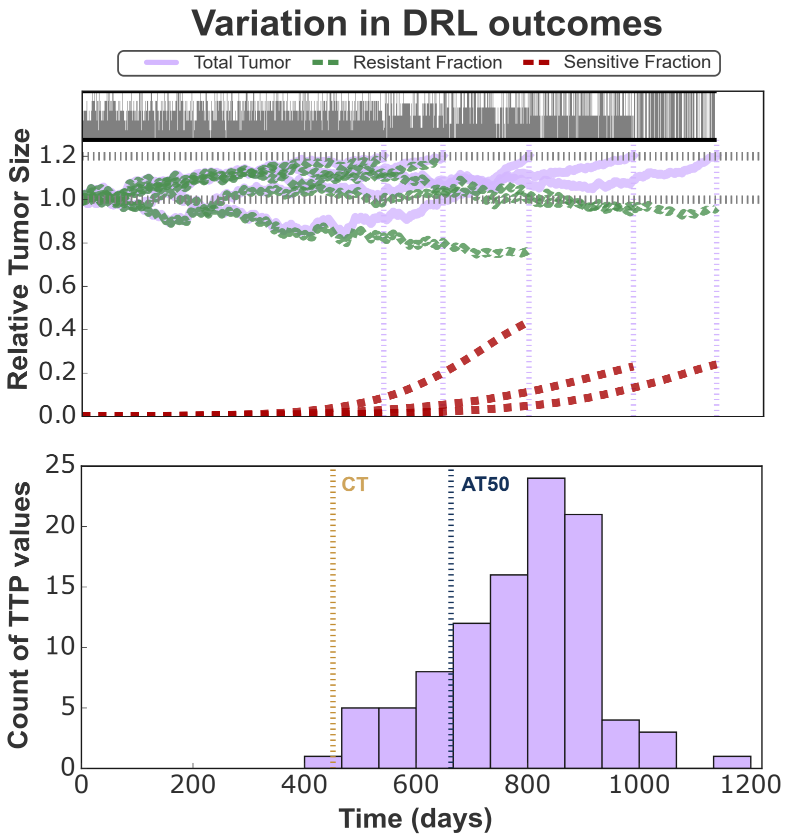


Figure 2: Variation in treatment outcomes due to stochasticity in the DRL's decision making process, evaluated on 100 copies of the virtual patient. 5 examples are shown in explicitly above, while the distribution of TTP for all 100 evaluations is depicted below, in comparison to the average TTP for continuous and adaptive therapy.

##### 5.2 DRL Robustness to Model Variation

In Section 3.4 we consider the robustness of the DRL framework to variation in patient dynamics. Through this, we introduce a modified Lotka– Volterra model introduced by [16], to demonstrate that a pre-trained DRL network can adapt to changes in the underlying tumor dynamics. Explicitly, this model may be written (in non-dimensional form) as:

$$\begin{aligned}\frac{dS}{dt} &= r_S S\left[1-\left(\frac{S+\frac{R}{1+e^{\gamma t}}}{K_S}\right)^{\alpha}-d_S D\right],\\ \frac{dR}{dt} &= r_R R\left[1-\left(\frac{R+\frac{S}{1+e^{\gamma t}}}{K_R}\right)^{\alpha}-d_R D\right],\end{aligned} \tag{17}$$

where $S$ and $R$ are the sensitive and resistant cell sub-populations respectively, and $D$ is the drug concentration.

This has been parameterised according to Table 2, replicating values used by Lu et al. [16], and chosen to ensure that the profile in question does reach progression:

| Name | Description | Value |
|---|---|---|
| $r_S$ | Sensitive cell proliferation rate | $0.01365\,day^{-1}$ |
| $r_R$ | Resistant cell proliferation rate | $0.00825\,day^{-1}$ |
| $K_S$ | Carrying capacity for sensitive cells | 1.0 |
| $K_R$ | Carrying capacity for resistant cells | 0.25 |
| $d_S$ | Drug-induced sensitive cell killing | 2.3205 |
| $d_R$ | Drug-induced resistant cell killing | 1.3205 |
| $S_0$ | Initial sensitive cell fraction | 0.75 |
| $R_0$ | Initial resistant cell fraction | 0.01 |
| $\alpha$ | Growth scaling term | 1.0 |
| $\gamma$ | Relative competition | $0.27385\,day^{-1}$ |

Table 2: Parameter values used for the alternative virtual patient model, taken from Lu et al. [16].

### References


[1] K. Arulkumaran, M. P. Deisenroth, M. Brundage, and A. A. Bharath, "Deep reinforcement learning: A brief survey," *IEEE Signal Processing Magazine*, vol. 34, pp. 26–38, Nov. 2017.

[2] J. Peters and S. Schaal, "Natural actor-critic," *Neurocomputing*, vol. 71, pp. 1180–1190, Mar. 2008.

[3] D. Silver, G. Lever, N. Heess, T. Degris, D. Wierstra, and M. Riedmiller, "Deterministic policy gradient algorithms," in *Proceedings of the 31st International Conference on Machine Learning* (E. P. Xing and T. Jebara, eds.), vol. 32 of *Proceedings of Machine Learning Research*, (Bejing, China), pp. 387–395, PMLR, June 2014.

[4] R. H. Crites and A. G. Barto, "An actor/critic algorithm that is equivalent to q-learning," in *Proceedings of the 7th International Conference on Neural Information Processing Systems*, NIPS'94, (Cambridge, MA, USA), p. 401–408, MIT Press, 1994.

[5] T. Degris, P. M. Pilarski, and R. S. Sutton, "Model-free reinforcement learning with continuous action in practice," in *2012 American Control Conference (ACC)*, IEEE, June 2012.

[6] V. Mnih, A. P. Badia, M. Mirza, A. Graves, T. Lillicrap, T. Harley, D. Silver, and K. Kavukcuoglu, "Asynchronous methods for deep reinforcement learning," in *Proceedings of The 33rd International Conference on Machine Learning* (M. F. Balcan and K. Q. Weinberger, eds.), vol. 48 of *Proceedings of Machine Learning Research*, (New York, New York, USA), pp. 1928–1937, PMLR, June 2016.

[7] S. Hochreiter and J. Schmidhuber, "Long short-term memory," *Neural Computation*, vol. 9, pp. 1735–1780, Nov. 1997.

[8] J. S. Bridle, "Probabilistic interpretation of feedforward classification network outputs, with relationships to statistical pattern recognition," in *Neurocomputing*, pp. 227–236, Springer Berlin Heidelberg, 1990.

[9] M. A. Strobl, J. West, Y. Viossat, M. Damaghi, M. Robertson-Tessi, J. S. Brown, R. A. Gatenby, P. K. Maini, and A. R. Anderson, "Turnover modulates the need for a cost of resistance in adaptive therapy," *Cancer Research*, vol. 81, pp. 1135–1147, Feb. 2021.

[10] J. Zhang, J. J. Cunningham, J. S. Brown, and R. A. Gatenby, "Integrating evolutionary dynamics into treatment of metastatic castrate-resistant prostate cancer," *Nature Communications*, vol. 8, Nov. 2017.

[11] J. A. Gallaher, P. M. Enriquez-Navas, K. A. Luddy, R. A. Gatenby, and A. R. Anderson, "Spatial heterogeneity and evolutionary dynamics modulate time to recurrence in continuous and adaptive cancer therapies," *Cancer Research*, vol. 78, pp. 2127–2139, Jan. 2018.

[12] E. Malaise, N. Chavaudra, and M. Tubiana, "The relationship between growth rate, labelling index and histological type of human solid tumours," *European Journal of Cancer (1965)*, vol. 9, pp. 305–312, Apr. 1973.

[13] J. B. West, M. N. Dinh, J. S. Brown, J. Zhang, A. R. Anderson, and R. A. Gatenby, "Multidrug cancer therapy in metastatic castrate-resistant prostate cancer: An evolution-based strategy," *Clinical Cancer Research*, vol. 25, pp. 4413–4421, Apr. 2019.

[14] S. Prokopiou, E. G. Moros, J. Poleszczuk, J. Caudell, J. F. Torres-Roca, K. Latifi, J. K. Lee, R. Myerson, L. B. Harrison, and H. Enderling, "A proliferation saturation index to predict radiation response and personalize radiotherapy fractionation," *Radiation Oncology*, vol. 10, July 2015.

[15] C. Grassberger, D. McClatchy, C. Geng, S. C. Kamran, F. Fintelmann, Y. E. Maruvka, Z. Piotrowska, H. Willers, L. V. Sequist, A. N. Hata, and H. Paganetti, "Patient-specific tumor growth trajectories determine persistent and resistant cancer cell populations during treatment with targeted therapies," *Cancer Research*, vol. 79, pp. 3776–3788, May 2019.

[16] Y. Lu, Q. Chu, Z. Li, M. Wang, and Q. Zhang, "Deep reinforcement learning identifies personalized intermittent androgen deprivation therapy for prostate cancer," May 2022.
